# Early beta oscillations in multisensory association areas underlie crossmodal performance enhancement

**DOI:** 10.1101/2021.05.26.445840

**Authors:** Georgios Michail, Daniel Senkowski, Martin Holtkamp, Bettina Wächter, Julian Keil

## Abstract

The combination of signals from different sensory modalities can enhance perception and facilitate behavioral responses. While previous research described crossmodal influences in a wide range of tasks, it remains unclear how such influences drive performance enhancements. In particular, the neural mechanisms underlying performance-relevant crossmodal influences, as well as the latency and spatial profile of such influences are not well understood. Here, we examined data from high-density electroencephalography (N = 30) and electrocorticography (N = 4) recordings to characterize the oscillatory signatures of crossmodal facilitation of response speed, as manifested in the speeding of visual responses by concurrent task-irrelevant auditory information. Using a data-driven analysis approach, we found that individual gains in response speed correlated with reduced beta power (13-25 Hz) in the audiovisual compared with the visual condition, starting within 80 ms after stimulus onset in multisensory association and secondary visual areas. In addition, the electrocorticography data revealed a beta power suppression in audiovisual compared with visual trials in the superior temporal gyrus (STG). Our data suggest that the crossmodal facilitation of response speed is associated with early beta power in multisensory association and secondary visual areas, presumably reflecting the enhancement of early sensory processing through selective attention. This finding furthers our understanding of the neural correlates underlying crossmodal response speed facilitation and highlights the critical role of beta oscillations in mediating behaviorally relevant audiovisual processing.

**Significance Statement:** The use of complementary information across multiple senses can enhance perception. Previous research established a central role of neuronal oscillations in multisensory perception, but it remains poorly understood how they relate to multisensory performance enhancement. To address this question, we recorded electrophysiological signals from scalp and intracranial electrodes (implanted for presurgical monitoring) in response to simple visual and audiovisual stimuli. We then associated the difference in oscillatory power between the two conditions with the speeding of responses in the audiovisual trials. We demonstrate, that the crossmodal facilitation of response speed is associated with beta power in multisensory association areas during early stages of sensory processing. This finding highlights the importance of beta oscillations in mediating behaviorally relevant audiovisual processing.

## Introduction

In everyday life, using complementary information from multiple sensory modalities is often critical to make rapid and accurate perceptual decisions. The synthesis of signals from different senses has been shown to improve perceptual performance, leading to more accurate (Spence and Driver, 2004; Lippert et al., 2007) and faster responses (Hershenson, 1962; Diederich and Colonius, 2004). Previous research has shown that crossmodal interactions are governed by neural oscillations in different frequency bands that can occur at both early and late stages of processing and involve bottom-up and top-down mechanisms (Keil and Senkowski, 2018; Bauer et al., 2020). Despite the considerable progress in characterizing the role of neural oscillations in multisensory processing, it remains unclear how they relate to the behavioral facilitation of responses to multisensory stimuli. In particular, the processing stage at which functionally relevant oscillations unfold during crossmodal behavior facilitation, and whether they reflect top-down or bottom-up influences on sensory processing, are key questions that are not well understood (Bizley et al., 2016).

In relation to the crossmodal facilitation of response times (RTs), electrophysiological studies in humans examining multisensory interactions in evoked brain potentials have suggested a link of RT facilitation with early crossmodal interactions (Giard and Peronnet, 1999; Fort et al., 2002; Molholm et al., 2004; Gondan et al., 2005). However, the proposed association in these studies is based on activity differences between multisensory and unisensory conditions that were not directly linked with the individual gains in multisensory performance enhancement. Thus far, only few studies have examined how neural oscillations relate to crossmodal RT facilitation across individuals (Senkowski et al., 2006; Mercier et al., 2015). In a speeded response paradigm, Senkowski et al. (2006) found a relationship between evoked beta oscillations and shorter RTs for unisensory and bisensory audiovisual stimuli. In an electrocorticography (ECoG) study, Mercier et al. (2015) observed that delta band (<4 Hz) phase alignment in a sensorimotor network was related to crossmodal facilitation of response speed. However, in both studies the modulations in neural oscillations were associated with shorter RTs after both multisensory and unisensory stimulation. Therefore, it cannot be concluded that these brain responses are specific for crossmodal facilitation of RTs. Moreover, the use of speeded responses in these studies, with a mean RT lower than 300 ms for audiovisual trials, indicates that the observed oscillatory activities may reflect motor-related processing. Taken together, while there is some evidence that neural oscillations play a role in crossmodal facilitation of response speed, the specificity of these effects to multisensory processing has not yet been demonstrated. Critically, it remains unclear whether the crossmodal facilitation of response speed is associated with modulations of neural oscillations during early stages of sensory processing.

In two experiments, we examined how individual gains in response speed during crossmodal stimulation relate to neural processing, as reflected in neural oscillations. We investigated oscillatory power in response to unisensory visual and bisensory audiovisual stimuli in experiments in which participants had to indicate the number of perceived flashes. Electrophysiological data were collected independently in healthy individuals (N = 30) using high-density EEG recordings and in patients with drug-resistant focal epilepsy (N = 4) prior to resective surgery, using ECoG recordings. The EEG data analysis revealed that lower early beta band power for audiovisual compared with visual trials in multisensory association and secondary visual regions correlated with crossmodal facilitation in response speed. The ECoG data analysis revealed lower beta power in audiovisual compared with visual trials in the superior temporal gyrus (STG). Our findings suggest that early beta band power in multisensory association cortex plays an important role in crossmodal facilitation of response speed.

## Material and Methods

The electrophysiological data from high-density scalp EEG and intracranial ECoG recordings were obtained independently. Throughout the text, the recording sessions to obtain these data are referred to as ‘EEG experiment’ and ‘ECoG experiment’, respectively.

### Participants

For the EEG experiment, forty participants (mean age ± standard deviation (SD): 26.6 ± 7.8 years; 19 females) with normal hearing, normal or corrected-to-normal vision and no history of neurological disorders were recruited. Six participants with excessive EEG artefacts (slow wave drifts and muscular artefacts) and four with insufficient trials (less than 30 trials in at least one of the analyzed conditions) were excluded from the analysis. Therefore, a subset of thirty participants (mean age ± SD: 25.5 ± 6.4 years; 17 females) was included in further EEG data analyses.

Four male patients (mean age ± SD: 27.3 ± 4.9 years) with drug-resistant focal epilepsy treated at the Epilepsy-Center Berlin-Brandenburg (Institute for Diagnostic of Epilepsy) in Berlin participated in the ECoG experiment. The patients were implanted with subdural electrodes (n = 66, 50, 40 and 74 for patients 1 to 4, respectively) covering mainly the temporal cortex for presurgical intracranial video-EEG monitoring.

All participants provided written informed consent. The experiments were conducted in accordance with the 2008 Declaration of Helsinki and approved by the ethics committee of the Charité–Universitätsmedizin Berlin (Approval number: EA1/207/15).

### Experimental Design

Participants were presented with combinations of auditory and visual stimuli and had to indicate the number of perceived visual stimuli. Stimulus combinations consisted of 0, 1 or 2 auditory (a) stimuli combined with either 0, 1 or 2 visual (v) stimuli. Six stimulus combinations were used in the EEG experiment, (a_0_v_1_, a_0_v_2_, a_1_v_1_, a_2_v_0_, a_2_v_1_, a_2_v_2_), and nine in the ECoG experiment (a_0_v_1_, a_0_v_2_, a_1_v_0_, a_1_v_1_, a_1_v_2_, a_2_v_0_, a_2_v_1_, a_2_v_2_ and a_2_v_1late_). The current study focused on the analysis of the visual-only stimulus (a_0_v_1_, **V**) and the bisensory audiovisual (a_1_v_1_, **AV**) stimulus combination in which one visual stimulus is presented together with one auditory stimulus (**Figure 1A**). In the EEG experiment, prior to the audiovisual stimulation, participants performed an n-back task (0-back, 2-back). In the current study, we only analyzed the a_0_v_1_ and a_1_v_1_ trials and only from the 0-back condition. Further details of the experimental setup can be found in Michail et al. (2021), which analyzed the memory-load effects on the perception of the a_2_v_1_ trials from the same EEG dataset. The visual (flash) stimulus was a white disk subtending a visual angle of 1.6° and was presented at 4.1° centrally below the fixation cross, for 13.3 ms (EEG) or 16.7 ms (ECoG). The slight difference in visual presentation times is explained by the different refresh rates of the displays used for the EEG and ECoG experiments. The auditory (beep) stimulus was a 78 dB (SPL) 1000 Hz sine wave tone that was presented for 7 ms. In AV trials, auditory and visual stimuli were presented simultaneously.

**Figure 1.**
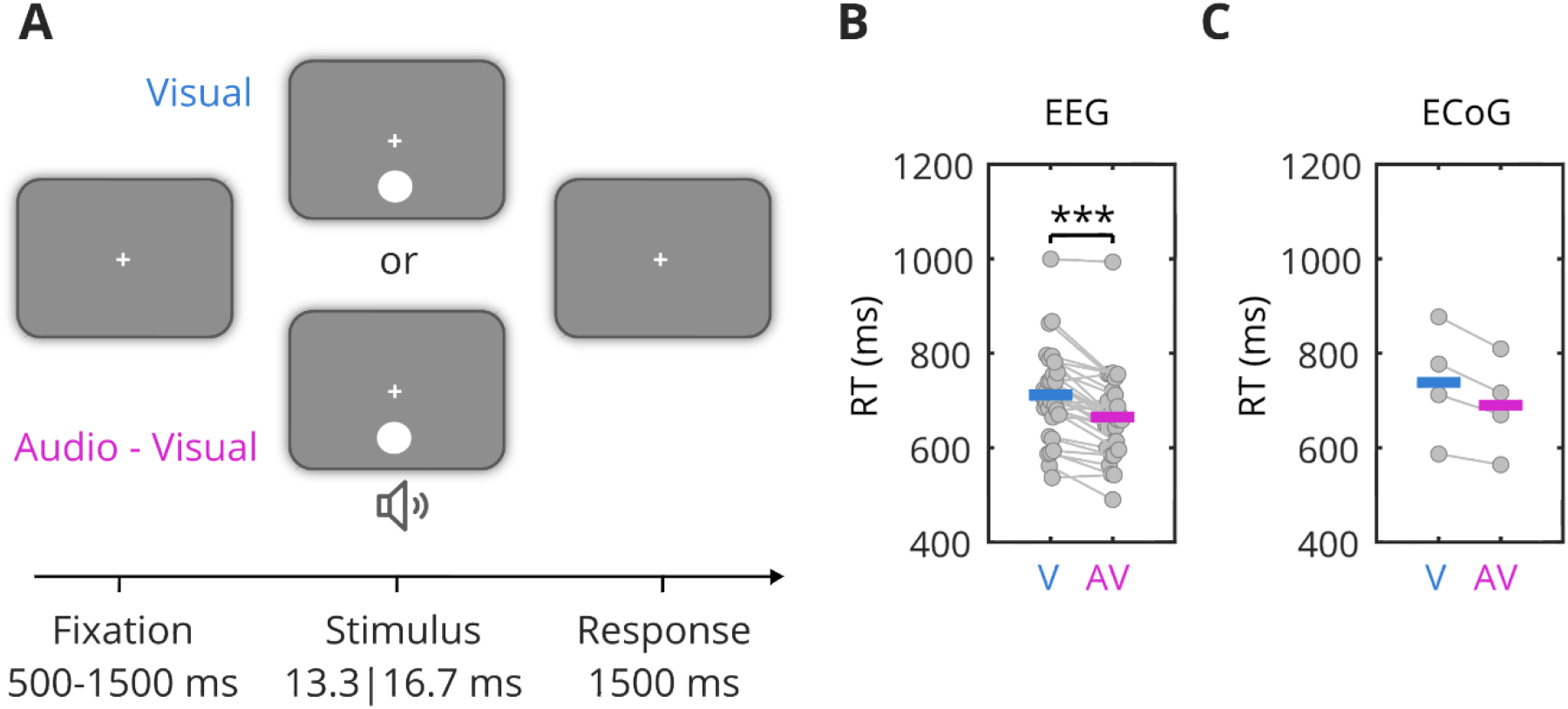
Experimental setup and behavioral results. **A.** Schematic illustration of the experimental conditions. Participants were presented with a unisensory visual (V) or a bisensory audio-visual stimulus (AV) and were asked to indicate the number of perceived visual stimuli. **B.** Participants in the EEG experiment responded faster in the AV compared with the V condition. Horizontal bold lines denote the mean. **C.** In the ECoG experiment, participants showed a speeding of responses in the AV condition, similar to the EEG experiment. Within-subject response speed was faster for AV compared with V stimuli in 3 out of 4 participants (significant or trend to significant difference). **\*\*\****p* < 0.001

Stimulus presentation and recording of participants’ responses were implemented using the Psychophysics toolbox (Brainard, 1997; RRID:SCR_002881) for MATLAB (The Mathworks, Natick, MA, USA). The EEG experiment was conducted in a dimly lit, electrically shielded, noise-attenuating chamber. Visual stimuli were displayed on a 21-inch CRT screen at a distance of 1.2 m with a 75 Hz refresh rate. The ECoG experiment was conducted at the patient’s bedside using a portable computer (HP Pavilion 17) with a 60 Hz screen refresh rate. Auditory stimuli in both experiments were controlled by a USB audio interface (UR22mkII, Steinberg) and delivered through in-ear headphones (ER30, Etymotic Research).

Each trial started with a central fixation cross displayed for a variable duration of 500 to 800 ms (EEG) or 1000 to 1500 ms (ECoG). Then, one of the stimulus combinations was presented. After the presentation of a stimulus, the fixation cross was displayed again and participants had to indicate the number of perceived flashes by a button press (three buttons: 0, 1 or 2). Following the button press or after 1500ms (if no button was pressed), a new trial started. In the EEG experiment, prior to the main task described above, participants performed a verbal visual n-back task (0- and 2-back, for details see Michail et al., 2021). In the current study, we only used trials from the 0-back condition, in which participants had to detect the target letter ‘X’, presented in 33% of all trials. The letter detection task was not related to the V and AV stimuli and should, thus, not have substantially affected the processing of these stimuli. Participants reported the number of flashes with the right thumb using a handheld gamepad (Logitech Gamepad F310, Logitech, Lausanne, Switzerland).

The EEG experiment included 12 blocks (6 blocks for each load level: 0-back and 2-back), each block consisting of 74 trials. The order of blocks was randomized across participants and the duration of experiment was approximately 80 minutes. The ECoG experiment, with a duration of 60 minutes, consisted of 6 blocks, each including 139 trials (due to fatigue, the first participant completed only 4 blocks).

### Behavioral data analysis

Behavioral performance was assessed in terms of the percentage of correct responses in the V and the AV condition and the RTs in trials with correct responses.

### Data acquisition and preprocessing

High-density EEG was recorded using a 128-channel passive system (EasyCap, Herrsching, Germany) at a sampling rate of 2500 Hz. Two electrodes, at the right lateral canthi and below the right eye, recorded the horizontal and vertical electro-oculograms. Preprocessing was performed with MNE-Python (Gramfort et al., 2014; RRID:SCR_005972) and further data analysis with Fieldtrip (Oostenveld et al., 2011; RRID:SCR_004849) and custom-made Matlab scripts (MathWorks, Natick, MA).

Offline, EEG data were filtered with a zero-phase bandpass finite impulse response (FIR) filter between 1 Hz and 100 Hz using the window design method (“firwin” in SciPy [https://docs.scipy.org/doc/]; Hanning window; 1 Hz lower transition bandwidth; 25 Hz upper transition bandwidth; 3.3 s filter length). A band-stop notch FIR filter from 49 to 51 Hz (6.6 s filter length), was applied to remove line noise. In the next analysis step, data were downsampled to 256 Hz and epoched from −1.5 to 1.5 s relative to the onset of the stimuli. Trials with artefacts (eye blinks, noise, or muscle activity) were removed after visual inspection. Data were then re-referenced to the average of all electrodes and subjected to Independent Component Analysis (ICA) using the Extended-Infomax algorithm (Lee et al., 1999). Components representing eye blinks, cardiac and muscle activity were removed from the data. Next, noisy electrodes were rejected after visual inspection on a trial-by-trial basis and interpolated using spherical spline interpolation (Perrin et al., 1989). Finally, trials with signal exceeding ±150 μV were excluded. On average, across participants, 106.5 (SD 96) trials and 12.1 (SD 4.3) ICA components were removed, and 11.1 (SD 3.6) electrodes were interpolated.

ECoG signals were recorded at a 2048 Hz sampling rate using a 128-channel REFA system (TMSi International, Enschede, The Netherlands). Offline, ECoG data were filtered using a zero-phase bandpass finite impulse response (FIR) filter between 1 Hz and 200 Hz (high pass: order = 6765, −6 dB cutoff frequency = 0.5 Hz; low pass: order = 137, −6 dB cutoff frequency = 225 Hz). A band-stop notch filter was applied at 50 Hz (±1) and its harmonics to filter out line noise. Data were subsequently downsampled to 600 Hz and epoched from −1 to 2.5 s relative to the onset of the stimulus. Electrodes with epileptiform activity or excessive noise were excluded from the analysis. Moreover, trials with an amplitude larger than five times the SD for more than a period of 25 ms (Blenkmann et al., 2019) and trials with artefacts (large slow drifts or excessive noise) identified after visual inspection were removed. Data were then re-referenced to the common average. On average, across participants in the ECoG experiment, 11.4 % (SD 4.3) of the trials and 7.4 % (SD 5.8) of the electrodes were removed.

To determine the locations of the intracranial electrodes, the post-implantation CT was co-registered with the preoperative MRI following the pipeline implemented in FieldTrip for the integrated analysis of anatomical and ECoG data (Stolk et al., 2018).

### Time-frequency analysis

Oscillatory power was computed by applying a Hanning taper to an adaptive time window of 4 cycles for each frequency from 2 to 40 Hz, shifted from −1.5 to 1.5 s (EEG) and from −1 to 2.5 s (ECoG), in steps of 10ms. Poststimulus power was baseline corrected using the average power of the prestimulus window from −500 to −100 ms, relative to stimulus onset.

### EEG source analysis

Surface-level EEG data were projected into source space to investigate the cortical sources of the correlation between spectral power and RTs, obtained from the sensor level analysis. First, for each participant, the individual T1-weighted MRI (3T Magnetom TIM Trio, Siemens, AG, Germany) was co-registered with the individually digitized EEG electrode positions (Polhemus FastTrak) to a common coordinate system (Montreal Neurological Institute, MNI). This was done by utilizing the digitized headshape information and the fiducial locations (nasion, left and right preauricular points). The co-registered MRI image was then segmented using the SPM12 algorithm and a realistic three-shell (brain, skull, skin) boundary element volume conductor model (BEM) was constructed (Oostendorp and van Oosterom, 1989). Next, the template MNI brain was non-linearly warped onto each participant’s anatomical data to obtain a three-dimensional source model (volumetric grid) with a resolution of 10 mm, which was used for the further analysis. To estimate the current density distribution the eLoreta algorithm (Pascual-Marqui, 2007) was used with a lambda regularization parameter set to 1%. To this end, the cross-spectral density (CSD) matrix was calculated using the Fast Fourier Transform (FFT) method for the condition-pooled data. As mentioned in the Introduction, in the current study, we were particularly interested on whether crossmodal RT facilitation is associated with early crossmodal influences. Accordingly, the source analysis focused on the early beta band component (80-200 ms, 13-25 Hz) of the significant cluster obtained from the scalp level correlation analysis. Therefore, CSD was calculated in the time window from 80 to 200 ms relative to stimulus onset. Center frequency and spectral smoothing were defined to fit the frequency range of interest; hence, a center frequency of 19 Hz and a smoothing of 6 Hz were used, resulting in a 13–25 Hz range. The current density estimate was normalized to the source estimate for the baseline window (−0.5 to −0.1 s) as follows: (*Poststimulus - Baseline*) / (*Poststimulus + Baseline*).

### Statistical analysis

For the EEG experiment, paired-samples *t-*tests were used to compare behavioral performance, i.e., accuracy and RTs, between V and AV conditions. The corresponding within-subject comparisons in the ECoG experiment were performed using independent-samples *t*-tests.

To compare the EEG spectral power between V and AV conditions, a nonparametric cluster-based permutation test was conducted (cluster-forming alpha = 0.05, dependent t-test, iterations = 1000; Maris and Oostenveld, 2007). The test was applied in the time window from 0 to 500 ms relative to stimulus onset, on frequencies from 2 to 40 Hz. The observed test statistic was evaluated against the permutation distribution in order to test the null hypothesis of no difference between conditions (two-tailed test, alpha = 0.025).

A nonparametric cluster-based permutation test was also applied to assess the correlation between the AV minus V power difference at the sensor level and the RT difference between the two conditions (cluster-forming alpha = 0.05, Spearman’s rank correlation, iterations = 1000). Accordingly, a similar approach was used for the corresponding correlation analysis of the source space data (one-sided cluster-based permutation test, cluster-forming alpha = 0.1, Spearman’s rank correlation, iterations = 1000). As mentioned before, the source analysis aimed to further investigate the findings of the sensor level analysis. Therefore, the direction of the one-tailed test was determined by the sensor level results.

With regard to the analysis of the ECoG data, the difference in beta power (averaged across the 13-25 Hz range) between V and AV conditions was assessed for each electrode in the time window from 0 to 500 ms using a nonparametric cluster-based permutation test (cluster-forming alpha = 0.05, independent samples *t*-test, iterations = 1000). Given that the non-symmetric arrangement of grid and strip electrodes prevents the use of spatial clustering algorithms, a more restricted alpha threshold of p = 0.01 was applied.

## Results

### Behavior

Behavior was assessed in terms of how fast and how accurate participants responded to V and AV stimuli. As depicted in **Figure 1B**, participants in the EEG experiment responded faster in the AV compared with the V condition (mean ± SD: 665 ± 92 ms vs. 712 ± 97 ms; paired samples *t*-test, t_(29)_ = 6.7, *p* < 0.001). Similarly, within-subject comparisons for participants in the ECoG experiment revealed significantly faster or a trend for faster responses in AV compared with the V condition in 3 out of 4 participants (**Figure 1C;** independent samples *t*-test, participant #1: 670 ± 97 ms vs. 712 ± 98 ms, t_(91)_ = −2.1, *p* = 0.038; #2: 716 ± 157 ms vs. 777 ± 193 ms, t_(150)_ = −2.2, *p* = 0.033; #3: 809 ± 182 ms vs. 877 ± 197 ms, t_(131)_ = −2.1, *p* = 0.041; #4: 564 ± 88 ms vs. 587 ± 87 ms, t_(150)_ = −1.6, *p* = 0.11). Only participant #4 revealed similar performance between conditions (p = 0.11). As this participant responded much faster than the other three participants, it is possible that the absence of a RT facilitation is due to a ceiling effect in performance. In the EEG experiment, while responses were more accurate in AV than V trials (98.1 ± 2.3 % vs. 92.0 ± 8.9 %, t(29) = −3.9, p < 0.001), participants showed in general high accuracy (>90%), suggesting that the task was easy to perform. Similarly, responses in the ECoG experiment were also highly accurate (V: 92.7 ± 5 %, AV: 93 ± 6.9 %; individual accuracies: participant #1: V=90.4%, AV= 88.5%; #2: V=96.2%, AV= 100%; #3: V=86.8%, AV= 85.9%; #4: V=97.4%, AV= 97.4%). Taken together, behavioral data from both EEG and ECoG experiments revealed that participants responded faster when the visual stimulus was combined with a task-irrelevant auditory stimulus than when the visual stimulus was presented alone.

### Audiovisual stimulation induces increased EEG theta power

In the first step we analyzed the difference in EEG oscillatory power between the AV and V condition in the window from 0 to 500 ms, on frequencies from 2 to 40 Hz, using only correct trials. As illustrated in **Figure 2**, the nonparametric cluster-based permutation test revealed stronger theta power increase in the AV compared with the V condition, over medio-frontal and occipital electrodes in the time window from 0 to 400 ms relative to stimulus onset (*p* = 0.003).

**Figure 2.**
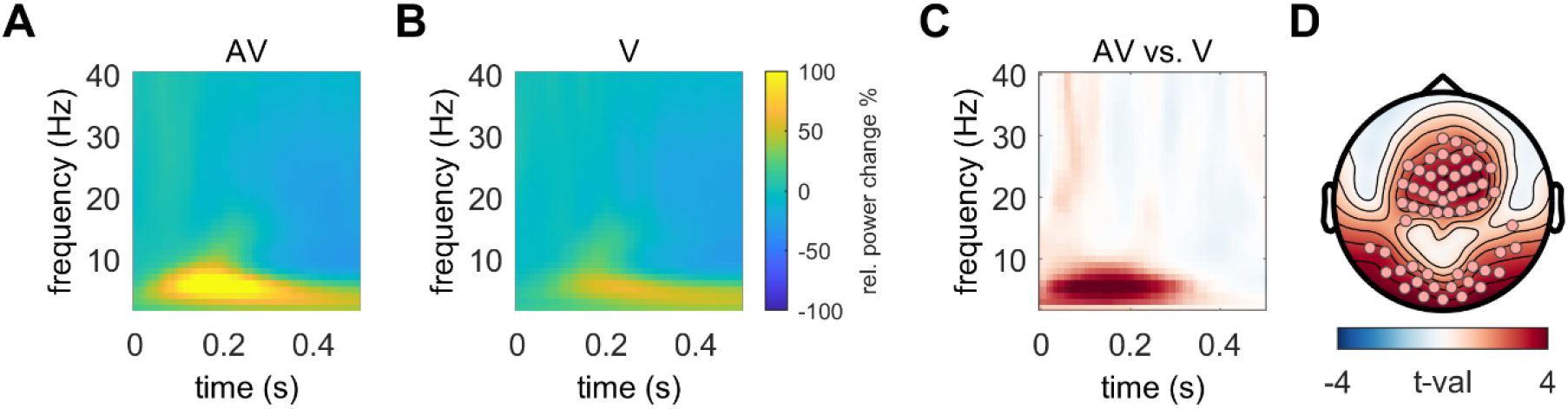
Oscillatory power difference between AV and V trials. The cluster-based analysis of EEG oscillatory power revealed higher theta power in AV compared with V trials in medio-frontal and occipital electrodes. **A-B.** TFRs of oscillatory power modulation after AV and V stimulation, averaged across electrodes with high contribution to the cluster (i.e., with a total number of significant time-frequency samples at or above the mean). **C.** TFR of AV-V power difference (in t-values), averaged across electrodes with high contribution to the cluster and masked based on the temporal and spectral extent of the cluster. Higher values indicate stronger power for AV compared with the V condition. The color scale refers only to unmasked t values. **D.** Topographic map showing the spatial distribution of the difference in the cluster’s time-frequency window. Electrodes with high contribution to the cluster are highlighted with dots.

### Early beta power in association areas correlates with crossmodal facilitation of response speed

We next examined whether differences in EEG oscillatory power between the AV and the V condition correlated with the crossmodal facilitation of RTs. (**Figure 1B**). For this analysis only correct trials were used. A nonparametric cluster-based permutation test revealed one significant cluster (*p* = 0.001) showing a negative correlation between the RT difference (Δ RT _V-AV_) and the power difference (Δ Power _AV-V_) over mainly parieto-occipital and frontal scalp regions (**Figure 3A-B**). The cluster comprised two components, one in the early beta band activity (strongest effect at 80-200 ms, 13-25 Hz) and a second one in the late alpha band activity (strongest effect at 250-400 ms, 8-12Hz). To confirm the finding of the cluster-based analysis, a Spearman’s rank correlation was performed between the RT facilitation (V minus AV) and the AV minus V power difference in the cluster (**Figure 3C;** rho = −0.81, *p* < 0.001). A comparison of the power in the cluster between the V and AV conditions revealed no significant difference between the two conditions. As mentioned in the *Introduction*, a central aim of the current study was to identify potential crossmodal effects at early processing stages. Therefore, the corresponding correlation analysis for the source activity focused on the early beta band activity (80-200 ms, 13-25 Hz). This analysis revealed a significant negative correlation of the AV minus V beta power difference in areas of the right inferior parietal and extrastriate occipital cortex with the crossmodal RT facilitation (nonparametric cluster-based permutation test, *p* = 0.001).

**Figure 3.**
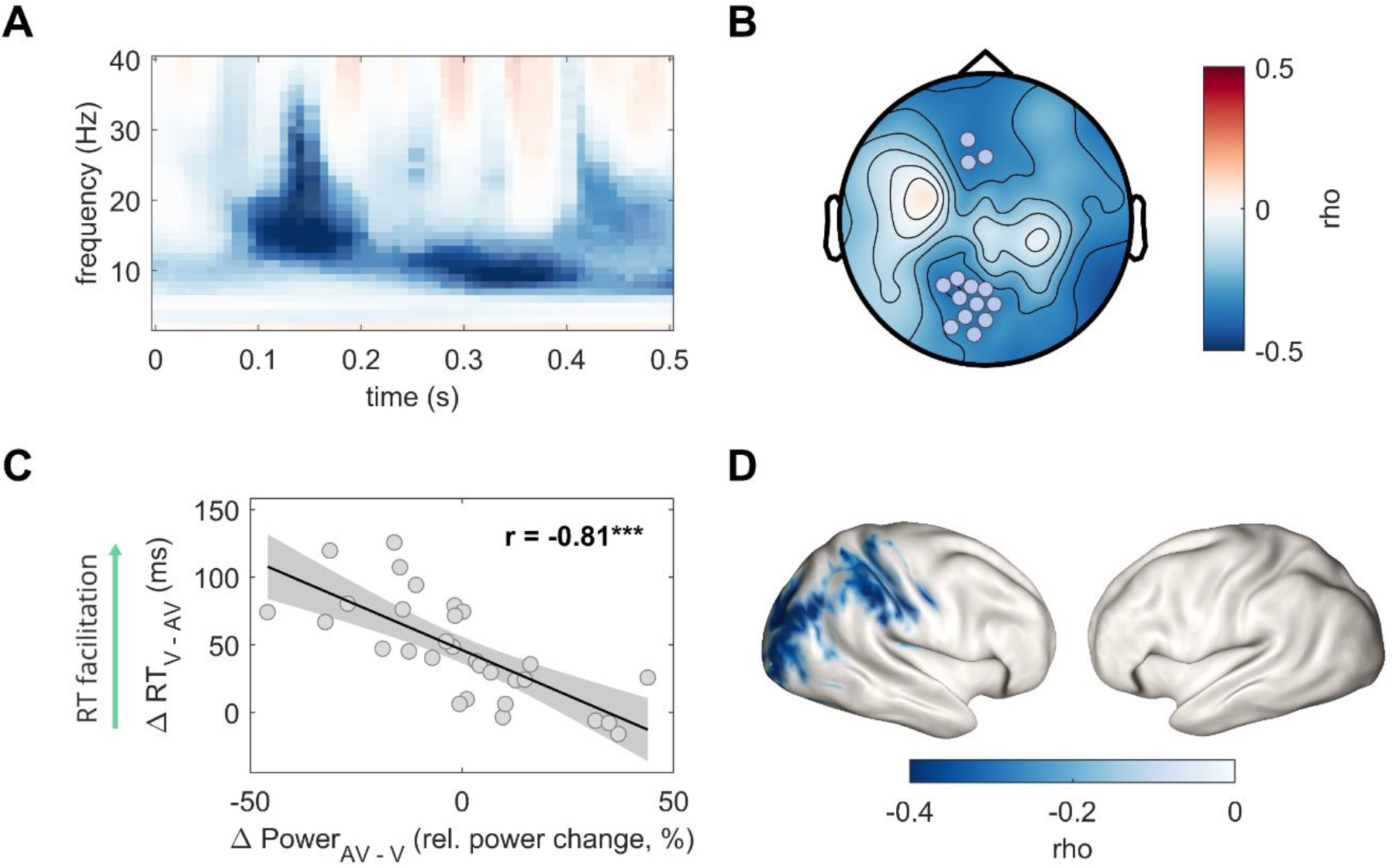
Correlation between AV minus V power difference and the crossmodal RT facilitation. The cluster-based correlation analysis revealed that crossmodal RT facilitation was associated with reduced beta power at 80-200 ms and reduced alpha power at 250-400 ms in mainly parieto-occipital electrodes, with the earlier beta effect being localized in inferior parietal and extrastriate occipital areas. **A.** TFR of the correlation (in rho values) between the AV minus V power difference and the V minus AV RT difference, averaged across electrodes with the highest contribution to the cluster (i.e., with a total number of significant time-frequency samples at or above the 75th percentile) and masked based on the temporal and spectral extent of the cluster. Lower values (blue) indicate that crossmodal RT facilitation correlates with smaller AV minus V power difference. The color scale refers only to unmasked rho values. **B.** Topographic map showing the distribution of the correlation between AV minus V power difference and the crossmodal RT facilitation. Electrodes with the highest contribution to the cluster are highlighted with dots. **C.** Scatterplot depicting the correlation between the individual power difference (A minus V) in the cluster and the crossmodal RT facilitation (i.e., V minus AV). The lower the power in the cluster for the AV compared with the V condition the larger the crossmodal RT facilitation. Black lines represent the best-fitting linear regression and shaded areas the 95% confidence interval. **D.** Correlation in source space between the early beta band power difference (AV minus V, 80-200 ms, 13-25 Hz) and the crossmodal RT facilitation. Lower AV vs. V beta power in inferior parietal and extrastriate visual areas correlated with the crossmodal RT facilitation.; **\*\*\****p* < 0.001

Taken together, our analysis revealed that the lower the early, parieto-occipital beta power in the AV compared with the V condition the faster participants responded in the AV vs. V condition. Moreover, the source localization of this correlation suggests the involvement of multisensory association areas and secondary visual cortex during the crossmodal RT facilitation.

### ECoG beta power in the superior temporal gyrus is lower in audiovisual compared with visual-only trials

To further examine the role of beta power during crossmodal processing, we compared beta power modulations between the unisensory V and bisensory AV conditions in four participants implanted with intracranial electrodes covering mainly the temporal cortex (**Figure 4**). As reported above, these participants displayed shorter RTs for the AV compared with the V condition (**Figure 1C**). Our primary interest in this study, was to investigate early crossmodal influences on neural oscillations. Therefore, based on the outcome of the EEG data analysis – which linked early beta power modulations with crossmodal RT facilitation – the ECoG data analysis focused on the time course of the beta band power (13-25 Hz). This analysis revealed that, consistently for all four participants, beta power in the superior temporal gyrus (STG) starting at approximately 130 to 150 ms poststimulus was significantly lower in the AV compared with the V condition (nonparametric cluster-based permutation test, *p* < 0.01). Interestingly, the reverse pattern was observed for very early beta power (< 100 ms) in few electrodes in participant #1 (STG) and participant #4 (rolandic operculum, middle frontal gyrus). In these electrodes, beta power in the first 100 ms after stimulus onset was significantly higher in AV compared with V trials. **Table 1** provides an overview of the statistical results and the MNI coordinates of the electrodes at which significant effects were observed. These results provide further evidence that beta band power modulations in multisensory association areas, and especially in STG, reflect early crossmodal influences that might play a critical role in crossmodal RT facilitation.

**Figure 4.**
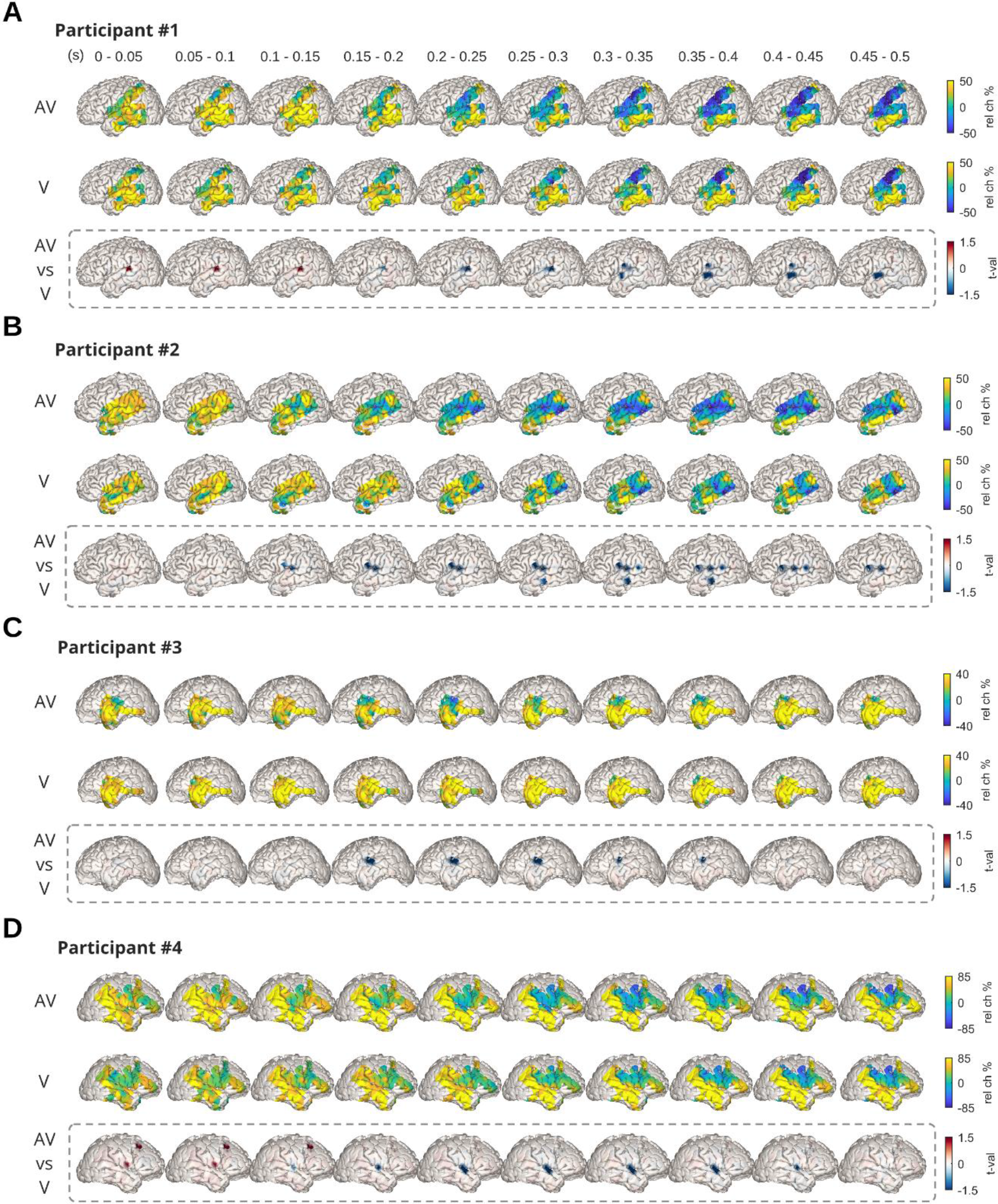
Intracranial (ECoG) beta band power in response to AV and V stimuli. The comparison of ECoG beta band power between the AV and the V condition showed that, consistent across participants, beta power in STG starting at approximately 150 ms after stimulus onset was significantly lower in AV compared with the V condition. **A-D.** For each participant, the first two rows display the beta band (13-25 Hz) power modulation after AV and V stimulation in the time window from 0 to 500 ms after stimulus onset. The third row (highlighted with a dotted line) shows the beta band power difference (in *t*-values) between AV and V conditions. Larger values (red) indicate stronger power for AV compared with the V condition.

**Table 1.**
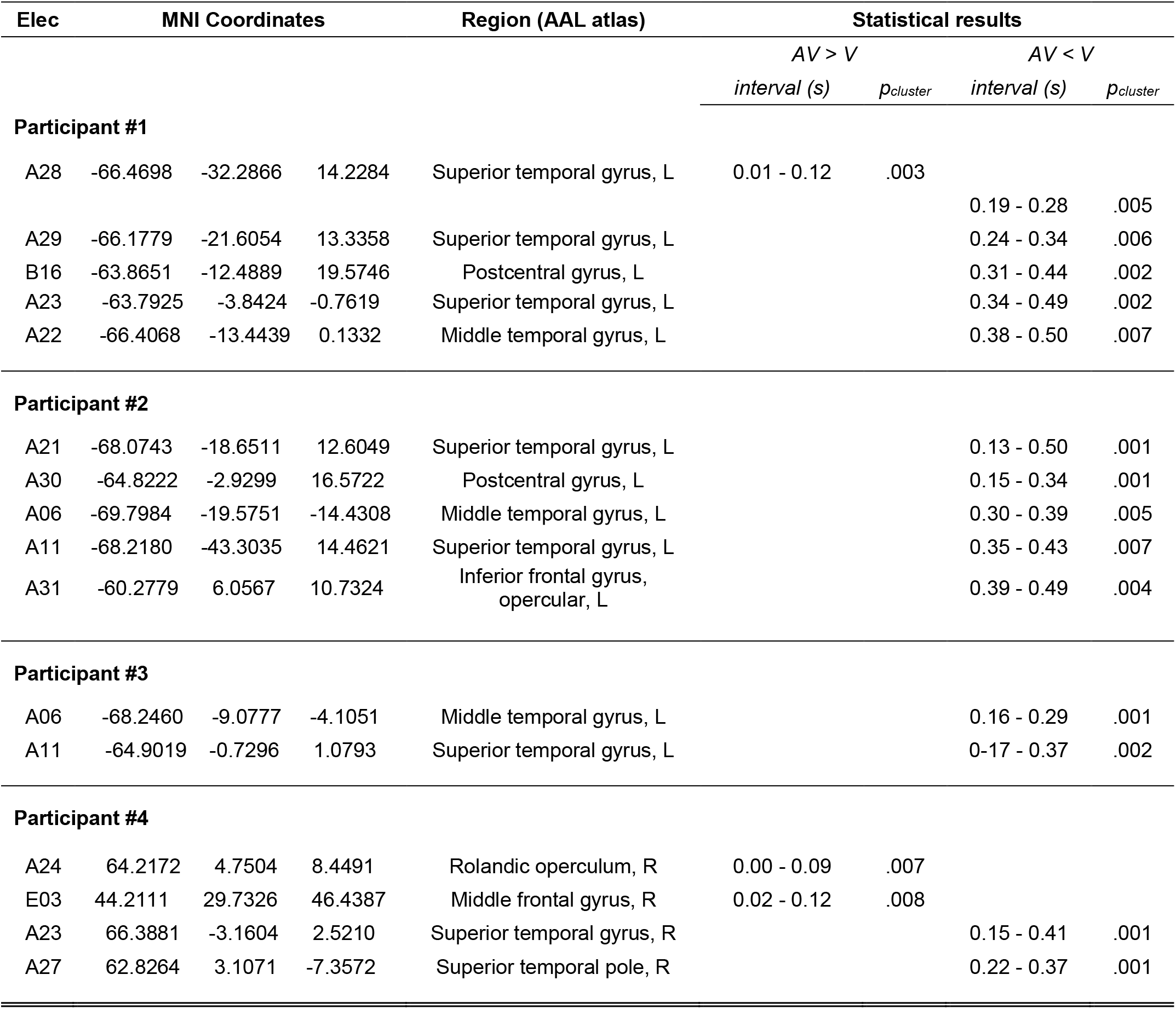
Comparison of beta power between V and AV trials in ECoG experiment

| Elec | MNI Coordinates |  |  | Region (AAL atlas) | Statistical results |  |  |  |
| --- | --- | --- | --- | --- | --- | --- | --- | --- |
|  |  |  |  |  | AV > V |  | AV < V |  |
|  |  |  |  |  | *interval (s)* | *p<sub>cluster</sub>* | *interval (s)* | *p<sub>cluster</sub>* |
| **Participant #1** |  |  |  |  |  |  |  |  |
| A28 | -66.4698 | -32.2866 | 14.2284 | Superior temporal gyrus, L | 0.01 - 0.12 | .003 |  |  |
|  |  |  |  |  |  |  | 0.19 - 0.28 | .005 |
| A29 | -66.1779 | -21.6054 | 13.3358 | Superior temporal gyrus, L |  |  | 0.24 - 0.34 | .006 |
| B16 | -63.8651 | -12.4889 | 19.5746 | Postcentral gyrus, L |  |  | 0.31 - 0.44 | .002 |
| A23 | -63.7925 | -3.8424 | -0.7619 | Superior temporal gyrus, L |  |  | 0.34 - 0.49 | .002 |
| A22 | -66.4068 | -13.4439 | 0.1332 | Middle temporal gyrus, L |  |  | 0.38 - 0.50 | .007 |
| **Participant #2** |  |  |  |  |  |  |  |  |
| A21 | -68.0743 | -18.6511 | 12.6049 | Superior temporal gyrus, L |  |  | 0.13 - 0.50 | .001 |
| A30 | -64.8222 | -2.9299 | 16.5722 | Postcentral gyrus, L |  |  | 0.15 - 0.34 | .001 |
| A06 | -69.7984 | -19.5751 | -14.4308 | Middle temporal gyrus, L |  |  | 0.30 - 0.39 | .005 |
| A11 | -68.2180 | -43.3035 | 14.4621 | Superior temporal gyrus, L |  |  | 0.35 - 0.43 | .007 |
| A31 | -60.2779 | 6.0567 | 10.7324 | Inferior frontal gyrus,<br>opercular, L |  |  | 0.39 - 0.49 | .004 |
| **Participant #3** |  |  |  |  |  |  |  |  |
| A06 | -68.2460 | -9.0777 | -4.1051 | Middle temporal gyrus, L |  |  | 0.16 - 0.29 | .001 |
| A11 | -64.9019 | -0.7296 | 1.0793 | Superior temporal gyrus, L |  |  | 0.17 - 0.37 | .002 |
| **Participant #4** |  |  |  |  |  |  |  |  |
| A24 | 64.2172 | 4.7504 | 8.4491 | Rolandic operculum, R | 0.00 - 0.09 | .007 |  |  |
| E03 | 44.2111 | 29.7326 | 46.4387 | Middle frontal gyrus, R | 0.02 - 0.12 | .008 |  |  |
| A23 | 66.3881 | -3.1604 | 2.5210 | Superior temporal gyrus, R |  |  | 0.15 - 0.41 | .001 |
| A27 | 62.8264 | 3.1071 | -7.3572 | Superior temporal pole, R |  |  | 0.22 - 0.37 | .001 |

## Discussion

In this study, we analyzed EEG and ECoG data to elucidate the neural correlates of the crossmodal RT facilitation, as manifested in the speeding of behavioral responses to visual stimuli by the addition of task-irrelevant auditory information. We showed that reduced beta power in the AV compared with V trials correlated with individual crossmodal RT gains. This effect occurred around 80-200 ms poststimulus in parieto-occipital electrodes and was localized in secondary visual and multisensory association areas. Moreover, the ECoG data analysis showed that beta power in the STG, which is a key multisensory association area, is reduced in AV compared with V trials, starting approximately 150 ms after stimulus onset. These findings provide evidence that beta band power modulations in multisensory association and secondary visual cortex during early visual sensory processing reflect the crossmodal facilitation of response speed.

Despite evidence of crossmodal influences occurring during both early and late multisensory processing and in both primary sensory and higher-order cortical areas (Macaluso and Driver, 2005; Koelewijn et al., 2010; Talsma et al., 2010; Keil and Senkowski, 2018), it is not well understood how such interactions enable the multisensory facilitation of performance. A central question regards the processing stage and the level of cortical hierarchy at which information from one modality influences another modality, in particular when such multisensory influences facilitate performance (Bizley et al., 2016).

Our finding that crossmodal RT facilitation was linked with oscillatory power modulations at 80-200 ms poststimulus suggests that the auditory signal influenced early visual sensory processing to enhance performance. This result is consistent with a large body of primate and human electrophysiological studies demonstrating multisensory interactions at early processing stages (Giard and Peronnet, 1999; Molholm et al., 2002; Schroeder and Foxe, 2005; Talsma and Woldorff, 2005; Lakatos et al., 2007; Kayser et al., 2010; Mercier et al., 2013). Moreover, our finding is in line with EEG studies providing direct evidence of early crossmodal responses underlying multisensory behavior facilitation in tasks using simple audiovisual stimuli (Thorne et al., 2011; Van der Burg et al., 2011; Starke et al., 2020). On the contrary, other studies using more complex stimuli have shown that sound-induced improvements of visual motion and visual object categorization were associated with late single-trial EEG activity starting at 300 ms (Kayser et al., 2017; Franzen et al., 2020). This divergence in the latency of performance-relevant crossmodal influences is consistent with evidence of multisensory integration taking place during both sensory encoding and decision formation (Mercier and Cappe, 2020) and is likely attributed to stimulus complexity, in accordance with the adaptive engagement of integrative mechanisms depending on task-specific characteristics (van Atteveldt et al., 2014; Bizley et al., 2016). In this framework, our data argue that under conditions of low stimulus complexity, multisensory RT facilitation is linked with crossmodal influences at early processing stages.

Critically, the crossmodal RT facilitation in our study was associated with power modulations in the beta band (13-25 Hz). The correlation between crossmodal beta power modulation and RT facilitation was observed in parieto-occipital electrodes and was localized in inferior parietal and extrastriate occipital regions. We propose that the performance-relevant beta power suppression in the audiovisual compared with the visual condition reflects the enhancement of early visual processing through top-down attentional control originating from multisensory association and secondary visual cortex. This proposal is consistent with growing evidence on the role of beta oscillations in conveying feedback influences on low-level visual areas (Buschman and Miller, 2007; Kerkoerle et al., 2014; Bastos et al., 2015; Michalareas et al., 2016; Richter et al., 2017; Limanowski et al., 2020). Moreover, evidence of feedback influences in the alpha-beta band modulating feedforward gamma band processing (Spaak et al., 2012; Richter et al., 2017) suggests that feedback signals in the low-frequency range (i.e., in the alpha-beta range), originating from association areas can directly modulate the feedforward stream of information during early sensory processing (Bressler and Richter, 2015). Our proposal is further supported by research showing that the suppression of low-frequency activity is associated with more efficient sensory processing of task-relevant signals (Klimesch et al., 2007; Jensen and Mazaheri, 2010), possibly by enhancing the feedforward communication through gamma band coherence (Hahn et al., 2019). In multisensory settings, previous studies provided evidence implicating beta power in the audiovisual redundant target effect (Senkowski et al., 2006), the integration of incongruent or noisy audiovisual speech stimuli (Schepers et al., 2013; Roa Romero et al., 2015), crossmodal influence on pain (Senkowski et al., 2011; Mancini et al., 2013), and the impact of working memory load on audiovisual illusory perception (Michail et al., 2021). Moreover, previous research demonstrated the involvement of beta band functional connectivity between primary and higher-order association areas in multisensory perception (Kayser and Logothetis, 2009; Hipp et al., 2011; Keil et al., 2014). Interestingly, a crossmodal (AV minus V) theta power enhancement over medio-frontal and occipital regions was not related to performance enhancement, suggesting that crossmodal theta power modulations might not be directly relevant for behavior. In this context, we argue that the functionally relevant beta band suppression in secondary visual and multisensory association areas – driven by the task-irrelevant auditory stimulus – enhanced early sensory representations of the visual stimulus through top-down attentional control of feedforward information processing.

Additionally, the analysis of the ECoG data revealed that beta power was reduced in the STG in the AV compared with the V condition. Previous work has established the critical role of the STG in multisensory perception, acting as a convergence hub for inputs from multiple modalities (Calvert et al., 2000; Beauchamp et al., 2004; Barraclough et al., 2005; Balz et al., 2016; Ozker et al., 2017; Karas et al., 2019; Mégevand et al., 2020). Moreover, previous studies using illusory audiovisual paradigms demonstrated that beta band power suppression was associated with audiovisual mismatch evaluation and top-down influences on audiovisual integration, induced by working memory load (Roa Romero et al., 2015; Michail et al., 2021). In accordance with these studies, the beta band suppression in the STG might reflect an auditory-driven feedback signal to improve visual processing through selective attention. This notion is consistent with the temporal and spatial profile of the observed tight relationship between beta oscillations in the EEG data and the crossmodal RT facilitation. It also in line with neuroimaging and electrophysiological evidence showing anatomical and functional connections in the beta band between STG and primary sensory areas (Noesselt et al., 2007, 2010; Cappe et al., 2009; Kayser and Logothetis, 2009; Keil et al., 2014). Therefore, this finding, together with the sources of the correlation between EEG beta power and RT facilitation, suggest an important role of multisensory association areas during behaviorally relevant early crossmodal processing.

Thus far, only few studies have investigated the oscillatory signatures of crossmodal RT facilitation using similar audiovisual stimuli as the current study (Senkowski et al., 2006; Mercier et al., 2015). Contrary to present findings, one previous study found that the audiovisual RT facilitation was associated with increased evoked beta power in left frontal and right occipital electrodes (Senkowski et al., 2006). This inconsistent finding might be explained by differences in the task instructions. In the current study participants had to report on features of the visual stimulus, whereas in Senkowski et al. (2006) participants made speeded responses upon stimulus detection independent of modality. This resulted in markedly faster RTs in Senkowski et al. (2006) compared with the current study (mean RTs to AV trials: 255 ms vs. 665 ms, respectively). Thus, the beta modulations in that previous study were possibly related to motor processes, whereas, in the present study, there is an additional perceptual aspect. Using a similar speeded detection task, an ECoG study has linked crossmodal RT facilitation with local phase alignment and phase synchronization between auditory and motor cortex in the delta band (Mercier et al., 2015). The use of a speeded detection task in these studies makes it difficult to disentangle the oscillatory activities associated with audiovisual interactions in sensory and non-sensory stages of information processing. Further investigations are required to differentiate the contributions of beta power and functional connectivity at the level of sensory processing, decision-making and motor response.

One limitation of our study is the small sample size in the ECoG experiment, which prevented us from performing similar analyses as in the EEG experiment. In addition to that, the heterogeneity between participants in the cortical grid coverage, further constrained the ability to perform analyses across participants to obtain statistically robust results at the group level. Thus, future ECoG studies, recruiting larger participant cohorts and possibly with a more diverse cortical grid coverage could provide insights into the role of other regions in crossmodal performance enhancement.

Altogether, our data suggest that beta power in multisensory association areas is related to the crossmodal facilitation of response speed. This beta power modulation presumably reflects the earliest stage of behaviorally relevant audiovisual feedback processing in higher multisensory areas, starting around 80 ms after stimulus presentation. Thus, the present findings highlight the important role of beta oscillations in mediating behaviorally relevant crossmodal influences between the auditory and visual modalities.

